# Mitochondrial ACSS1 regulates the oncometabolite 2-hydroxyglutarate and *De Novo* Pyrimidine biosynthesis under nutrient-deprived conditions in lymphoma

**DOI:** 10.1101/2024.12.20.628231

**Authors:** Johnvesly Basappa, Aaron R. Goldman, Cosimo Lobello, Shengchun Wang, David Rushmore, Olga Melnikov, Neil V. Sen, Vinay S. Mallikarjuna, Priyanka Jain, Masoud Edalati, Kathy Q. Cai, Pin Lu, Reza Nejati, Hossein Borghaei, Kathryn E. Wellen, Mariusz A. Wasik

**Author notes:** **Corresponding author:** Johnvesly Basappa, Ph.D. Department of Pathology, Fox Chase Cancer Center, Philadelphia, PA 19111.

## Abstract

The mitochondrial Acetyl-CoA synthetase short-chain family member 1 (*ACSS1*) converts acetate, an energy source in nutrient-deprived conditions, to mitochondrial acetyl-CoA. However, the specific mechanism behind this process remains unknown. Here, we show that ACSS1 is overexpressed in patients with mantle cell lymphoma (MCL), diffuse large B cell lymphoma (DLBCL), chronic lymphocytic leukemia (CLL), and Ibrutinib (IBR)-resistant cell lines. The mitochondrial stress test showed reduced oxygen consumption in ACSS1 knockdown (KD) cell lines. ^13^C-acetate stable isotope tracing revealed that ACSS1 knockdown (KD) in MCL cell lines attenuates the flux of mitochondrial acetate to acetyl-CoA, acetylcarnitine, and TCA cycle intermediates, including glutamine and aspartate, which are precursors for de novo pyrimidine synthesis. Consistently, there was a decrease in the labeling of glutamate, aspartate, dihydroorotate, and orotate pools in KD cell lines. Pathway analysis revealed the enrichment of *de novo* pyrimidine synthesis metabolites and depleting ACSS1 impaired cell growth and potential vulnerability of IBR-resistant MCL cells. Further, we discovered that acetate is used to synthesize 2-hydroxyglutarate in an ACSS1-dependent manner, and it is involved in histone methylation. These results highlight a metabolic phenotype in MCL cells, showing their ability to metabolize acetate in nutrient-deprived conditions and provide new insights into the role of ACSS1 in cancer metabolism.

**Significance:** The enzyme ACSS1 catalyzes the conversion of acetate, an energy source in nutrient-deprived conditions, to mitochondrial acetyl-CoA. Our results show that ACSS1 may affect cancer metabolism, particularly in MCL. Studies using MCL cell lines showed that reducing ACSS1 activity affects acetate utilization for various metabolic pathways and influences the production of oncometabolite D-2-hydroxyglutarate. Additionally, reducing ACSS1 activity affects the expression of cyclin D1 and inhibits cell growth, highlighting the potential significance of ACSS1 in cancer metabolism.

## Introduction

Mantle cell lymphoma (MCL) is a highly aggressive type of B-cell non-Hodgkin lymphoma (NHL) and represents about 3–10% of all adult NHL cases in Western countries. The median overall survival is approximately 3-5 years from diagnosis^1^. MCL is characterized by a defining chromosomal translocation t(11;14)(q13;q32) involving the IgH/CCND1 locus, leading to overexpression of cyclin D1 gene and increased resistance to chemo-immunotherapy and tyrosine kinase inhibitors^2–5^. The overexpression of cyclin D alone does not cause MCL pathogenesis, but it promotes biosynthetic pathways, including glycolysis^6,7^, the pentose phosphate pathway, and nucleotide synthesis^8^. Acquired resistance to the Bruton tyrosine kinase (BTK) inhibitor ibrutinib (IBR), the first therapy approved by the FDA for MCL, is common and leads to poor clinical responses in patients. Metabolic reprogramming in MCL is associated with cancer cell growth, proliferation, metastasis, and therapeutic resistance. Continued research focusing on resistance mechanisms is needed to improve the efficacy of IBR therapy and prolong patient survival. Most metabolomics research efforts have focused on glucose metabolism due to the seminal observations of Warburg about the dominant role glucose plays in many basic biosynthetic processes^9^. In higher vertebrates, the rapid proliferation of cells is supported primarily by the consumption of glucose and glutamine^10^. Acetate is another nutritional source vital for the progression of many cancers, which commonly exhibit dysregulated metabolism, and it is especially important under nutrient-deprived conditions^11,15^. A recent study has shown that acetate promotes the expression of PD-L1 and immune evasion by upregulating c-Myc^12^. Indeed, acetate is also generated in significant quantities by the liver during ketogenic conditions such as prolonged fasting or diabetes and released into the bloodstream^13^. The mitochondrial Acetyl-CoA synthetase short-chain family member 1 (*ACSS1*) converts acetate, an energy source in nutrient-deprived conditions^11,14^, to mitochondrial acetyl-CoA. Mammals express three isoforms of the short-chain acetyl-CoA synthetases family (ACSS1, ACSS2, and ACSS3) that catalyze the conversion of acetate and coenzyme-A into acetyl-CoA^15,16^, which is crucial for cell growth and cell proliferation^17,18^. ACSS1 and ACSS3 localize in mitochondria, while ACSS2 is present in the nucleus and cytoplasm^19^. Among the three isoforms, ACSS2’s role in acetate metabolism and cancer progression has been well studied in various types of cancers, such as pancreatic cancer^20^, hepatocellular carcinoma^21,22^, glioblastoma^23^, breast cancer^24^, prostate cancer^24^, bladder cancer^25^ and cervical squamous cell carcinoma^26^. ACSS1 plays a pivotal role in thermogenesis during fasting, under low-glucose or ketogenic conditions, and is essential for survival^27^. Most acetyl-CoA is derived from glucose and glutamine pathways in highly oxygenated conditions. However, in hypoxic conditions, acetate is used for acetyl-CoA synthesis^28^. The previous paper outlined the function of ACSS1 in mitochondrial acetate metabolism, particularly in acute myeloid leukemia and melanoma^29^. The specific role of mitochondrial ACSS1 under nutrient-deprived conditions remains unknown. In this study, we investigate the role of ACSS1 in mitochondrial acetate metabolism using comprehensive ^13^C-glucose, ^13^C-glutamine, and ^13^C-acetate stable isotope tracing and ACSS1 knockdown approach. The expression of ACSS1 is regulated in an estrogen receptor-dependent manner, and the estrogen hormone plays a role in regulating the incorporation of acetate into DNA^30,31^. Here, we discovered a new role for ACSS1 under nutrient-deprived conditions, highlighting its role in acetate-mediated cell survival, including the synthesis of the oncometabolite 2-hydroxyglutarate, its effect on histone methylation, and the *de novo* pyrimidine synthesis.

## Materials and Methods

ACSS1 (Cat#17138-1-AP), ACLY (Cat# 67166-1-Ig), GAPDH (Cat# 60004-1-Ig) from Proteintech, ACSS2 (Cat#ab66038) from Abcam. Stable isotope labelled U-^13^C6-Glucose (Cat#CLM-1396-1), ^13^C5-L-Glutamine (Cat#CLM-1822-H) and ^13^C2-Sodium acetate (Cat#CLM-440-1) from Cambridge Isotope Laboratories. The special media, D-Glucose-free RPMI 1640 (REF 11879-020), was obtained from Gibco. RPMI 1640 (-) Glutamine free and (-) D-Glucose free media was obtained from BI, Biological Industries Israel Beit-Haemek.

### MCL, DLBCL, and ALCL cell lines

Cell lines from different lymphomas were used, including MCL-RL^32–34^, a cell line derived from a patient with MCL at the University of Pennsylvania, Philadelphia, PA, JeKo-1, Maver, Rec-1, and Granta519 for MCL; OCI-LY1, OCI-LY4, OCI-LY8, and TOLEDO for DLBCL; SUPM2, SR786 and SUDH-L1 for ALK+ ALCL. All commercially available cell lines were purchased from the ATCC. Moreover, in our study we employed a human CD4+ T-cells transduced with NPM-ALK cell line, namely NA1, created by our group as previously described^35^. All the cell lines were regularly tested for Mycoplasma contamination using Mycoplasma detection kits from Thermo Fisher Scientific. All cells were grown in RPMI medium supplemented with 10% FBS and 1% penicillin/streptomycin under a humidified 37°C/5% CO_2_ incubator.

### Nutrient starvation in cell culture

For nutrient-deprivation experiments, cells were washed once with phosphate-buffered saline. The glutamine-free medium was made with RPMI devoid of glutamine and glucose and retained the same nutrient concentrations as standard RPMI with dialyzed 10% FBS and 25 mM Glucose. A glucose-free medium was made with RPMI devoid of glutamine and glucose and retained the same nutrient concentrations as standard RPMI with dialyzed 10% FBS and 2 mM Glutamine. Complete RPMI was prepared by supplementing with 2 mM L-Glutamine and 25 mM Glucose to RPMI devoid of glutamine and glucose. For acetate supplementation, Sodium acetate at a final concentration of 5 mM was supplemented to Glutamine-free or Glucose-free media or both Glutamine/Glucose-free media.

### ACSS1 and mitochondrial fractionation

For mitochondria isolation, 50×10^6^ cells of Jeko-1 and Maver cell lines were harvested, and mitochondria were isolated according to the instructions of the product manual (catalog # 89874, Thermofisher). The protein estimation was measured, and an equal amount of protein was separated on a 10% NuPAGE gel and processed for western blotting as described below.

### ACSS1 knockdown (KD) in MCL cell lines

Jeko-1 and Maver cell lines were transduced with the scramble PLKO.1-PURO NON-TARGET CONTROL (SHC016V-1EA) and ACSS1 shRNA, TRCN0000436733, TRCN0000045380, TRCN0000045381, TRCN0000424197 and TRCN0000423168 all purchased from Millepore-Sigma-Aldrich. The ready-to-transduce lentivirus particles were directly mixed with 100,000 cells (Jeko-1 and Maver) in the presence of polybrene (catalog#TR-1003-G, Millepore Sigma) in a 96-well plate. The media was replaced with fresh RPMI after 16 hours of incubation. The cells were cultured for a further 3 days and selected for puromycin.

### Western blotting

Cells were harvested and processed for the western blotting as previously described^36^ with respective antibodies, blots were developed using the Ibright 1500 imaging system (Thermo Scientific).

### Immunohistochemistry (IHC) of MCL, DLBCL, and CLL patient-derived biopsy samples

Frozen DLBCL and MCL patient tissue samples were isolated from IRB-exempt discarded specimens of patients with diffuse large B-cell lymphoma (DLBCL) obtained through the Fox Chase Cancer Center Bio Specimen Repository. Human tissue microarrays for patients with MCL (*n* = 22), DLBCL (*n* = 28) and CLL (n=) were used to examine ACSS1. The expression of ACSS1 was analyzed according to the pathology grade of the samples. Tissues were collected and fixed in 10% phosphate-buffered formaldehyde (formalin) for 24-48 hrs, dehydrated and embedded in paraffin. Hematoxylin and eosin (H&E) stained sections were used for morphological evaluation and unstained sections for IHC studies. IHC staining was performed on a VENTANA Discovery XT automated staining instrument (Ventana Medical Systems) using VENTANA reagents according to the manufacturer’s instructions. Slides were de-paraffinized using EZ Prep solution (cat # 950–102) for 16 min at 72 °C. Epitope retrieval was accomplished with CC1 solution (cat # 950– 224) at high temperature (eg, 95–100 °C) for 32 min. Rabbit primary antibodies (ACSS1: 1:200, Rabbit, Proteintech, Cat. 17183-AP) were tittered with a TBS antibody diluent into user-fillable dispensers for use on the automated stainer. Immune complex was detected using the Ventana OmniMap anti-rabbit detection kit (760-4311) and developed using the VENTANA ChromMap DAB detection kit (cat # 760-159) according to the manufacturer’s instructions. Slides were then counterstained with hematoxylin II (cat # 790-2208) for 8 min, followed by Bluing reagent (cat # 760-2037) for 4 min. The slides were dehydrated with ethanol series, cleared in xylene, and mounted. As a negative control, the primary antibody was replaced with normal rabbit IgG to confirm the absence of specific staining. Stained slides were scanned using an Aperio ScanScope CS 5 slide scanner (Aperio, Vista, CA, USA). Scanned images were then viewed and captured with Aperio’s image viewer software (ImageScope, version 11.1.2.760, Aperio).

### Metabolomics analysis

Metabolomics analyses were conducted at The Wistar Institute Proteomics and Metabolomics Shared Resource with slight modifications as previously described^37^. For ^13^C-Glucose tracing experiments, the cell lines were cultured in glucose- and glutamine-free RPMI-1640 media supplemented with 10 mM uniformly labeled U-^13^C-glucose (Cambridge Isotope Laboratories Inc., D-Glucose-U^13^C_6_, 99%, CLM-1396) and 2 mM unlabeled Glutamine for 16 hours in triplicate. For ^13^C-Glutamine tracing experiments, the cells were cultured in glucose and glutamine-free RPMI-1640 media supplemented with 1 mM labeled ^13^C_5_-L-Glutamine (Cambridge Isotope Laboratories Inc.), 10 mM unlabeled Glucose for 16 hours in triplicate. For ^13^C_2_-acetate tracing experiments, the cells were cultured in glucose- and glutamine-free RPMI-1640 media supplemented with 500 µM labeled ^13^C_2_-acetate (Cambridge Isotope Laboratories Inc.), 10 mM unlabeled Glucose for 16 hours in triplicate. The cells were counted using the trypan blue method, and an equal number of cells (10×10^6^ cells/condition) were spun down at 1,000 rpm for 5 minutes. The cell pellet was washed with cold PBS and snap-frozen in liquid nitrogen. Unlabeled controls were generated for each cell line condition.

### Oxygen consumption rate (OCR) measurement by Seahorse analysis

The MCL cell lines (Jeko-1 shControl (control), Jeko-1 shACSS1 knockdown (KD), Maver shControl (control), Maver shACSS1 knockdown (KD) were exposed to either -glucose (Glc) or 2 mM acetate (ACE) for 1 hour and then thoroughly examined for mitochondrial respiration using the Seahorse-based method. The XF Mitochondrial stress test kit protocols (Agilent) were followed using the Agilent Seahorse XFe96 analyzer per the manufacturer’s instructions. In brief, cells were seeded in Seahorse XF 96-well plates at a density of 1.2x 10^5^ cells per well with Assay Base media, supplemented with 10 mM Glucose, 1 mM Sodium Pyruvate, and 2 mM Glutamine. The cell cultures were allowed to equilibrate for 1 hour at 37 °C in a no-CO2 incubator. The Oxygen Consumption Rate was analyzed under basal conditions and after treatment with different drugs, including 1 µM oligomycin A, 2 µM FCCP, and 0.5 µM Rot/AA. After the analysis, the medium was removed, cells were suspended in RIPA lysis buffer, and the protein content of cell lysates was measured by Bradford assay and used to normalize respiratory parameters. Samples were analyzed with at least 6 technical replicates. The data were assessed using XF Wave Software (Seahorse Bioscience, Agilent).

### Statistics and reproducibility

The student’s unpaired t-test was used to analyze differences in RNAseq, quantitative (q)-PCR expression analysis and isotope-labeled metabolites compared with RL cell line vs. Jeko-1 and Maver in three ^13^C-isotope tracing experiments. For ACSS1 knockdown (KD) studies in Jeko-1 and Maver cell lines, an unpaired t-test was used to analyze differences between control vs. KD cells ^13^C-acetate isotope-labeled metabolites. *P* values equal to or less than 0.05 were considered statistically significant without being adjusted for multiple comparisons.

### Data availability

We confirm that all relevant data and methods are included in the main Article and the Supplementary Information section. The source data for the graphs and charts in the figures is available as a Supplementary. The analysis results in Extended Data Fig. 1a and b are based upon data generated by the Expression Atlas, and Figure 1c is based upon data generated by the TCGA Research Network: http://cancergenome.nih.gov/. Data supporting this study’s findings are available within the article and from the corresponding author upon request.

**Fig. 1.**
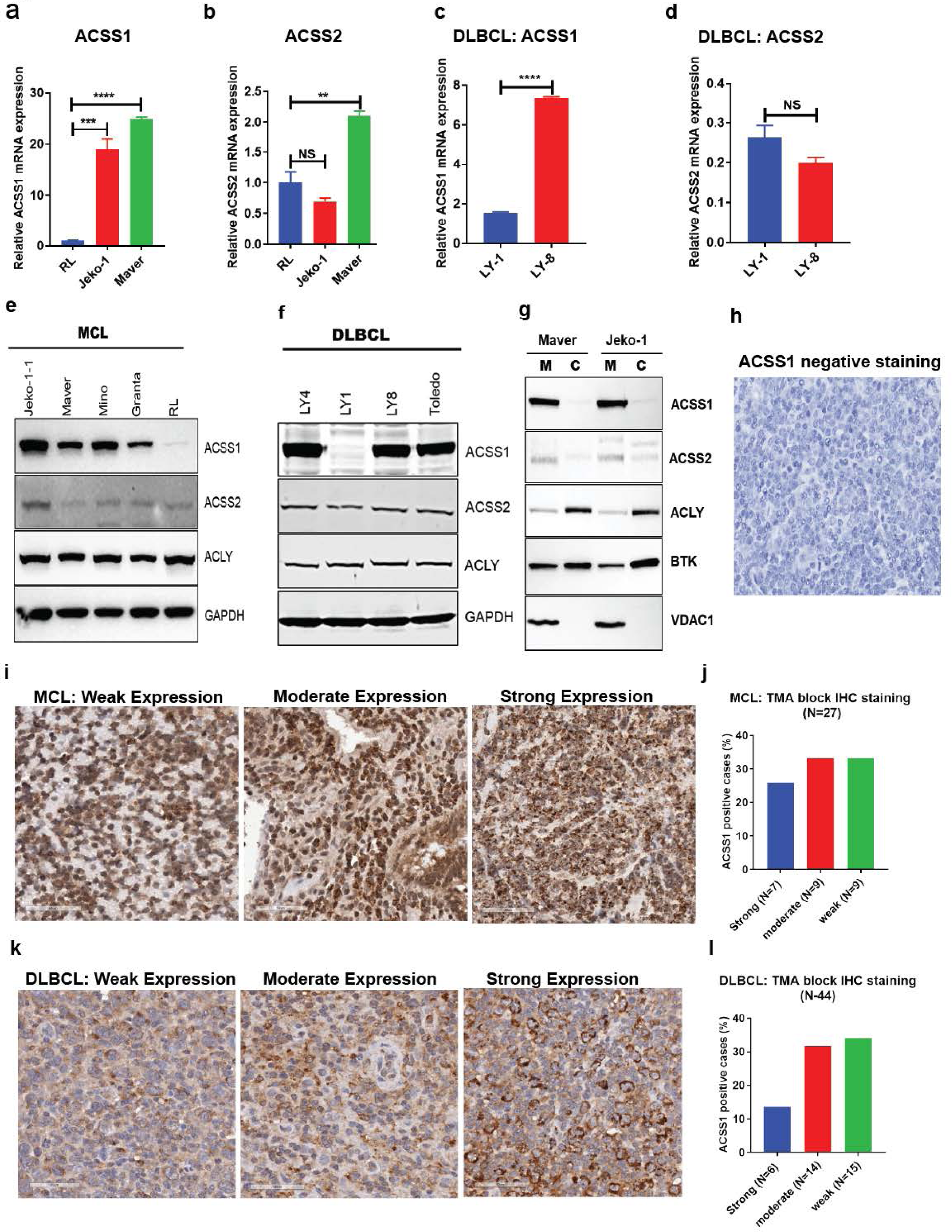
Mitochondrial ACSS1 is overexpressed in MCL, DLBCL, and ALCL cell lines. Q-PCR analysis of RL (sensitive to the BTK inhibitor ibrutinib), Jeko-1, and Maver (ibrutinib-resistant) MCL cell lines revealed increased expression of key acetate metabolic enzymes. **a-b**, ACSS1, and ACSS2 expression in MCL cell lines. **c-d**, ACSS1, and ACSS2 expression in DLBCL cell lines. **e**, ACSS1, ACSS2, and ACLY protein expression in MCL cell lines. **f,** ACSS1, ACSS2, and ACLY protein expression in DLBCL cell lines. **g**, Mitochondrial/cytosol fractionation of Maver and Jeko-1 cell lines. **h,** Negative Immunohistochemistry (IHC) staining of ACSS1 expression in MCL patient biopsy samples. **i,** IHC staining of ACSS1, weak expression (left), moderate expression (middle), and intense expression (right). (Representative image of n=27) **j**, The quantification of MCL patients based on their ACSS1 expression category. **k**, ACSS1 overexpression in DLBCL patient biopsy samples, weak expression (left), moderate expression (middle), and strong expression (right). (Representative image of n=44). **l**, The quantification of DLBCL patients based on their ACSS1 expression category. All graphs show mean ± SD (n = 2 biological replicates for q-PCR), and all statistical analyses were performed using PrismGraphPad for unpaired t-tests: ^∗^p < 0.05, ^∗∗^p < 0.01, ^∗∗∗^p < 0.001, ^∗∗∗∗^p < 0.0001.

## Results

### Mitochondrial ACSS1 is overexpressed in hematological malignancies and solid tumors

Acetyl-CoA synthetases are critical in converting acetate to acetyl-CoA through an ATP-dependent reaction. To gain insights into the potential impact of acetate metabolism and Acetyl-CoA synthetases in cancer models, we analyzed publicly available RNA-seq data from cell lines available in Expression Atlas (https://www.ebi.ac.uk/gxa/home) with a focus on hematological malignancies, emphasizing hematopoietic and lymphoid-derived cancer cell lines such as mantle cell lymphoma (MCL), diffuse large B-cell lymphoma (DLBCL), chronic lymphocytic leukemia (CLL), Acute Myeloid Leukemia (AML) and anaplastic large cell lymphoma (ALCL). This analysis revealed that mitochondrial ACSS1 is highly expressed in non-Hodgkin lymphoma (NHL) cancer cell lines and cancer patient samples (Extended Data Fig. 1a, 1b). Moreover, we observed a higher expression of ACSS1 than ACSS2 and ACSS3 in MCL cell lines. Additionally, we found that ACSS2, located in the cytosolic and nuclear compartments, is highly expressed in solid tumor cell lines (Extended Data Fig. 1c). In contrast, expression of ACSS3 across all the profiled cell lines was minimal (FPKM values 0 or <1) (Extended Data Fig. 1a, 1b). Next, we delved into the significance of ACSS1 and ACSS2 association with hematopoietic and lymphoid cancer patients, especially those with AML and DLBCL, using The Cancer Genome Atlas (TCGA) database^38^. ACSS1 showed high expression in samples derived from hematopoietic and lymphoid tissues, with 36.2% of the 221 tested samples exhibiting over-expression, while only 1.36% showed under-expression (Table 1). Next, we observed low expression of ACSS2 in lymphoid samples when compared to solid tumors (data not shown). Due to ACSS1 being understudied across all cancers, we also profiled ACSS1 expression patterns in solid tumors using TCGA data. We found that it is overexpressed in the following cancers relative to normal tissue: lung cancer (10.89%), ovarian cancer (29.32%), prostate cancer (16.47%), central nervous system tumors (79.91%), cervical cancer (84.04%), large intestine cancer (25.9%), and head and neck squamous cell carcinoma (24.71%) (Table 1).

**Table 1.** TCGA data revealed overexpression of ACSS1 in hematopoietic, lymphoid, and solid tumors.

| Cancer source/type | Total number of samples tested | No. of sample over expressed | Over expressed (%) | No of samples under expressed | Under expressed (%) |
| --- | --- | --- | --- | --- | --- |
| Hematopoietic and lymphoid | 221 | 80 | 36.2 | 3 | 1.36 |
| Central Nervous System | 697 | 557 | 79.91 | 32 | 4.59 |
| Lung | 1019 | 111 | 10.89 | None | None |
| Ovary | 266 | 78 | 29.32 | None | None |
| Cervix | 307 | 258 | 84.04 | 29 | 9.45 |
| Large intestine | 610 | 158 | 25.9 | 12 | 1.97 |
| Prostate | 498 | 82 | 16.47 | 3 | 0.6 |
| Head and Neck | 522 | 129 | 24.71 | None | None |

### ACSS1 is highly expressed in MCL, DLBCL, and ALCL cell lines

We conducted a detailed analysis of ACSS1 gene expression across multiple cell lines, including MCL and DLBCL. Our investigation revealed that ACSS1 expression was relatively low in the MCL-RL (RL) cells, which are known to be sensitive to Ibrutinib (IBR). In contrast, we identified two IBR-resistant MCL cell lines, Jeko-1 and Maver, that exhibited significantly higher levels of ACSS1 expression. Specifically, Jeko-1 demonstrated an impressive 18-fold increase (P < 0.0001), while Maver showed a remarkable 24-fold increase (P < 0.001) in comparison to the IBR-sensitive RL cell line (Fig.1a). But the expression of ACSS2 is not significant (Fig. 1b). The findings in DLBCL cell lines reveal an interesting pattern: ACSS1 is significantly more expressed in the OCI-LY8 cell line, which is resistant to IBR, compared to the IBR-sensitive OCI-LY1 cell line (Fig. 1c). Additionally, it is noteworthy that there was no significant expression of ACSS2 observed (Fig. 1d). These results suggest a potential relationship between ACSS1 expression and IBR resistance that warrants further investigation. Furthermore, we conducted immunoblotting on five MCL cell lines (Jeko-1, Maver, Mino, Granta, and RL). The results showed that ACSS1 is highly expressed in all the cell lines except the RL line, which is IBR sensitive (Fig. 1e). Based on these findings, we hypothesized that the expression of ACSS1 is associated with BTK inhibitor IBR sensitivity and resistance mechanism. Next, we immunoblotted four DLBCL cell lines (OCI-LY1, OCI-LY4, OCI-LY8, and Toledo), and the results showed high ACSS1 expression in all of the lines except the IBR-sensitive OCI-LY1 cell line (Fig. 1f). Based on TCGA data, we observed high expression of ACSS1 in hematopoietic cell lines. This observation was further validated in B-cell lymphoma (MCL and DLBCL). Our findings conclude that ACSS1 expression is higher in MCL and DLBCL cell lines less sensitive to IBR. To confirm the expression of ACSS1 in mitochondria, we conducted mitochondrial and cytosolic cell fractionation on MCL cell lines Maver and Jeko-1 following the protocol described in the methods. The fractionated protein lysates were then subjected to immunoblotting. The results revealed that ACSS1 expression is primarily located in the mitochondria (Fig. 1g). Based on the ACSS1 gene and protein expression pattern in MCL, DLBCL, and ALK+ALCL cell lines, we validated the extent of ACSS1 protein expression in MCL patient-derived biopsy samples (N=27) by conducting immunohistochemical (IHC) staining on tumor tissue microarrays available at the Fox Chase Cancer Center hospital. The negative staining of ACSS1 IHC is shown (Fig. 1h). The IHC data showed significant ACSS1 expression (N=25 with 92.59% total positivity) compared to normal tissue (Fig. 1i). The expression was categorized as strong (N=7, 25.93% positivity), moderate (N=9, 33.33% positivity), and weak (N=9, 33.33% positivity) in MCL patient samples (Fig. 1j). The complete TMA block in low magnification showed (Extended Data Fig.2a). Similarly, ACSS1 IHC staining was performed on DLBCL patient-derived tumor tissue microarrays (N=44). The results showed that ACSS1 total positivity in 35 patients (Fig. 1k). The expression was stratified into strong (N=6, 13.63% positivity), moderate (N=14, 31.82% positivity), and weak (N=15, 34.09% positivity) in DLBCL patient samples (Fig. 1l). The entire TMA block IHC results showed in lower magnification (Extended Data Fig.2 b). Next, IHC analysis of CLL (N=13) patient-derived tumor microarray biopsy samples showed ACSS1 overexpression. Among the thirteen CLL patient samples subjected to IHC staining, we observed (N=7) positivity with 53.84% (Extended Data Fig.2 a). Next, we created a tissue microarray (TMA) block derived from MCL, DLBCL, and ALCL cell lines. To assess ACSS1 expression, we conducted IHC staining on cell lines TMA using ACSS1 antibody. The results showed that the overexpression of ACSS1 in MCL cell lines, such as Jeko-1, Rec-1, and Granta in consistent with immunoblot and IHC analysis of patient-derived biopsy samples (Extended Data Fig.3a). Furthermore, the higher magnification images showed the overexpression of ACSS1 in MCL cell lines (top panel), DLBCL cell lines (middle panel), and ALCL cell lines (bottom panel) (Extended Data Fig.3b).

## 13C-labeled stable isotope tracing revealed Acetate fueling the TCA cycle in high ACSS1 expressing MCL cell lines

To investigate the correlation between ACSS1 expression and acetate metabolism, we performed stable isotope tracing analysis using ^13^C-labeled glucose, glutamine, and acetate and liquid chromatography-mass spectrometry (LC-MS) to calculate mass isotopologue distributions (MIDs) of TCA cycle intermediates and gain insight into the carbon source utilization of MCL-RL (RL) cells with low ACSS1 expression compared to Jeko-1 and Maver cell lines, which overexpress ACSS1. The isotopic tracers ^13^C-glucose, ^13^C-glutamine, and ^13^C-acetate create different labeling patterns in downstream metabolites involved in central carbon metabolism. This allows for comparison of substrate utilization in MCL cell lines with low and high expression of ACSS1, as shown in the schematics (Figure 2a, b, c). The labeling of M+2 Acetyl-CoA derived from ^13^C-glucose was consistent across the cell lines and showed no significant difference. (Fig. 2d). Similarly, the labeling of Acetyl-CoA from ^13^C-glutamine was 32% in RL, 33% in Jeko-1, and 41% in Maver (Fig. 2e). When comparing the labeling from ^13^C-glucose, there was no significant difference in ^13^C-glutamine tracing between low and high ACSS1 expressing cell lines (Fig. 2e). Next, we examined the M+2 labeled acetyl-CoA derived from ^13^C-acetate in RL, Jeko-1, and Maver cell lines. The results revealed a highly significant difference in the M+2 labeled Acetyl-CoA in RL vs. Jeko-1 (P < 0.0001) and RL vs. Maver ( P < 0.0001) in the ^13^C-acetate MIDs analysis (Fig 2f). Next, we focused on other TCA cycle intermediates citrate, α-KG, succinate, and malate incorporation of carbons derived from ^13^C-glucose,^13^C-glutamine, and ^13^C-acetate sources on M+2, M+4, and M+5 in multiple turns of the TCA cycle in three cell lines. The data revealed a mild difference in M+2 citrate labeling among three cell lines cultured in ^13^C-glucose media but nothing significant in ^13^C-glutamine (Fig. 2g-h). However, the results showed a significant difference in the citrate carbon labeling pattern (M+2; >22, 30 and 45%; p< 0.0007), M+4; >3, 21 and 17%; p<0.0001) and M+5, >1, 20 and 9%; p<0.0001) in RL, Jeko-1 and Maver cell lines respectively cultured in ^13^C-acetate media (Fig. 2i). Similarly, the data revealed that there was no difference in M+2, α-KG labeling among three cell lines cultured in ^13^C-glucose and ^13^C-glutamine (Fig.2j-k). The results showed the most significant difference in α-KG the carbon labeling pattern (M+2, M+3, M+4, and M+5) in Jeko-1 and Maver cell lines compared to RL cell line cultured in ^13^C-acetate (Fig. 2l). Next, we assessed the labeling of succinate and data revealed that there was no significant difference in M+2-n, succinate labeling among three cell lines cultured in ^13^C-glucose and ^13^C-glutamine (Fig.2m-n). Next, the data revealed a significant difference in succinate the carbon labeling pattern (M+2, M+3, M+4, and M+5) in Jeko-1 and Maver cell lines compared to RL cell lines cultured in ^13^C-acetate (Fig. 2o). Next, we observed similar results on malate M+2-n labeling pattern in ^13^C-glucose and ^13^C-glutamine (Fig.2p-q) and a huge significant difference in malate carbon labeling pattern (M+2, M+3, M+4, and M+5) in Jeko-1 and Maver compared to RL cell line cultured in ^13^C-acetate (Fig. 2r). Our carbon tracing experiments indicate that high expression of ACSS1 drives acetate metabolism in MCL cell lines, particularly under nutrient-deprived conditions.

**Fig. 2.**
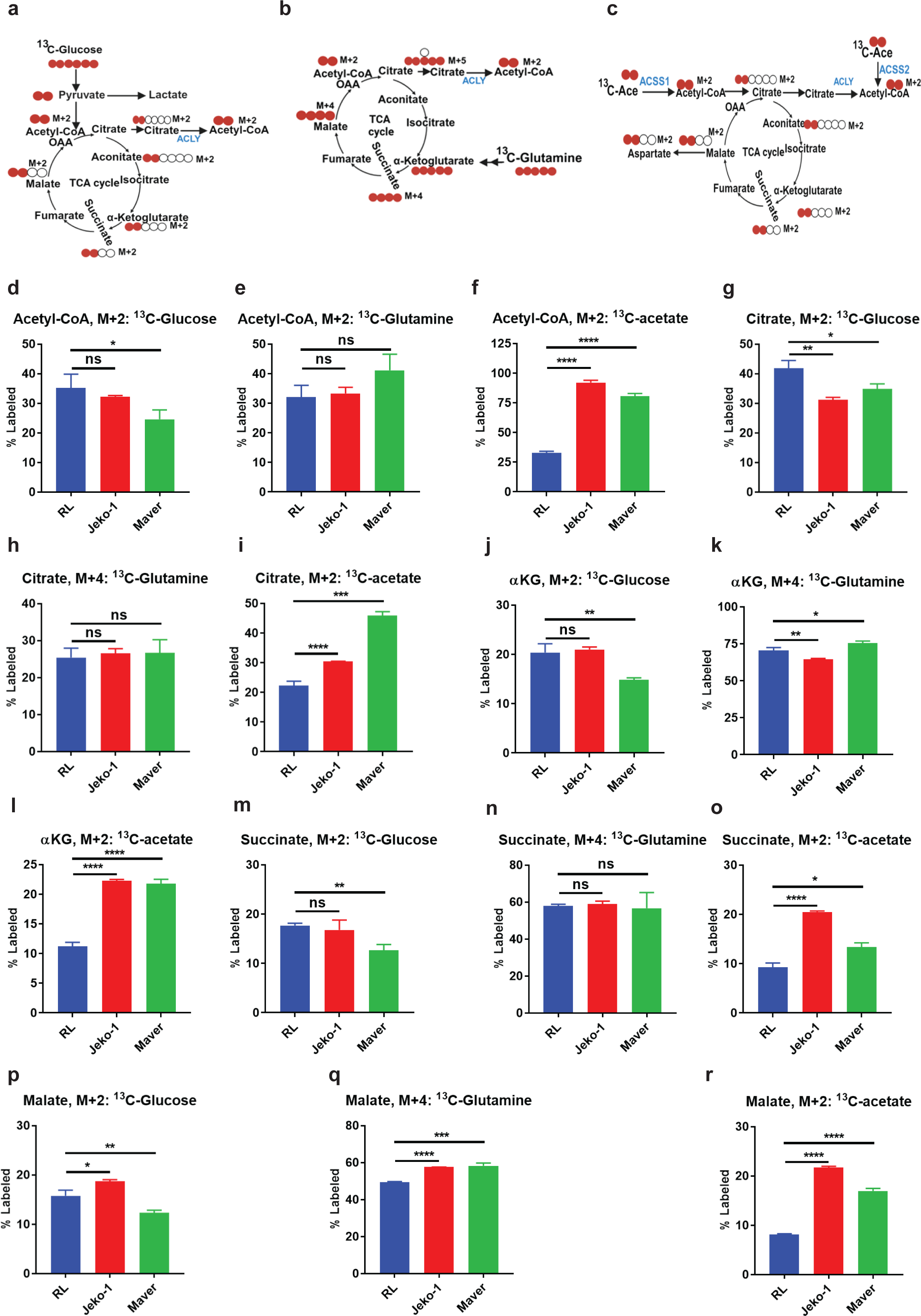
13C-isotopes tracing and mass isotopologues distributions (MIDs) in mantle cell lymphoma cell lines revealed increased acetate metabolism. **a**, A schematic of labeled carbon molecules from ^13^C-glucose is in red, and unlabeled ones are in black circles. **b,** A schematic of labeled carbon molecules from ^13^C-glutamine is in red, and unlabeled ones are in black circles. **c,** A schematic of labeled carbon molecules from ^13^C-acetate is in red, and unlabeled ones are in black circles. **d-f,** M+2 labeled acetyl-CoA from ^13^C-glucose, ^13^C-glutamine and ^13^C-acetate respectively. **g-i,** M+2-6 labeled citrate from ^13^C-glucose, ^13^C-glutamine and ^13^C-acetate respectively. **j-l,** M+2-5 labeled α-ketoglutarate (α-KG) from ^13^C-glucose, ^13^C-glutamine and ^13^C-acetate respectively. **m-o**, M+2-4 labeled succinate from ^13^C-glucose, ^13^C-glutamine and ^13^C-acetate respectively. **p-r**, M+2-4 labeled malate from ^13^C-glucose, ^13^C-glutamine and ^13^C-acetate respectively. All graphs show mean ± SD (n = 3 biological replicates), and all statistical analyses were conducted with unpaired t-tests: ^∗^p < 0.05, ^∗∗^p < 0.01, ^∗∗∗^p < 0.001, ^∗∗∗∗^p < 0.0001.

### ACSS1 silencing revealed a critical role in mitochondrial acetate metabolism

To better understand the role of ACSS1 in mitochondrial acetate metabolism, we silenced ACSS1 in two MCL cell lines, Maver and Jeko-1. The efficiency of ACSS1 knockdown was confirmed through q-PCR and immunoblotting (Fig. 3a, b). To test if ACSS1 is a crucial driver in mitochondrial acetate metabolism, we performed ^13^C-acetate flux analysis on both the shControl (control) and shACSS1 knockdown (KD) Jeko-1 and Maver cell lines as shown in schematic (Fig. 3c). The cells were cultured in ^13^C-acetate (1.0 mM) in glucose/glutamine-free RPMI media and supplemented with 1% dialyzed FBS for 24 hours. Metabolomics analysis results showed a significant reduction (>64%; P<0.0001) in the incorporation of Acetyl-CoA M+2 isotopologue in control *vs.* KD Jeko-1 cell lines (Fig. 3d), although no significant change was revealed in Maver cell line (Fig.3e). Subsequently, we tested whether the observed differences may be due to cytosolic or mitochondrial acetyl-CoA pools using mitochondrial acetylcarnitine as a readout^39–41^. The results showed that there was a significant reduction in M+2 carbon labeling in acetylcarnitine between the control and KD in Jeko-1 (>40%; P<0.0003) and Maver cell lines (>61%; P<0.0005), suggesting that ACSS1 is critical for the production of mitochondrial acetyl-CoA from acetate in both cell lines (Fig. 3f). Next, we compared the ^13^C-acetate derived mass isotopologue distribution (MIDs) in TCA cycle intermediates and gain insight into the role of ACSS1 in acetate utilization in the mitochondria of the Jeko-1 and Maver control and KD cell lines. The ^13^C-acetate tracing data revealed that ACSS1 KD resulted in a significant decrease in the labeling of citrate (M+2-5) that is generated over multiple rounds of the TCA cycle in Jeko-1 and Maver cell lines (Fig.3 g, h). Further investigation of other TCA cycle intermediates, including aconitate (Fig. 3i, j), α-KG (Fig. 3k, l), succinate (Fig. 3m, n), and malate (Fig. 3o, p), consistently showed significantly decreased proportions of labeled isotopologues derived from acetate in ACSS1 KD cell lines. These findings indicate that ACSS1 plays a critical role in acetate metabolism to power OXPHOS through the TCA cycle in MCL cell lines with high ACSS1 expression. As indicated, malate was highly labeled via the TCA cycle in the ^13^C-acetate tracing assay in control Jeko-1 and Maver cells (Fig. 3o, p). This process notably impacted the shACSS1 KD cell lines. Other studies have reported the participation of acetate in gluconeogenesis in hepatic cells using ^14^C-labeled radioisotopes^42^. Here, we show that ACSS1 metabolizes ^13^C-acetate to pyruvate and lactate in Jeko-1 and Maver cells, and there was reduced production of pyruvate and lactate derived from acetate in cell lines with ACSS1 knockdown (Fig.3q,r). This indicates that under nutrient-deprived conditions, MCL cells with high levels of ACSS1 convert acetate into energy through a series of reactions in the TCA cycle and synthesize glucose from pyruvate via gluconeogenesis, using non-carbohydrate precursors.

**Fig. 3.**
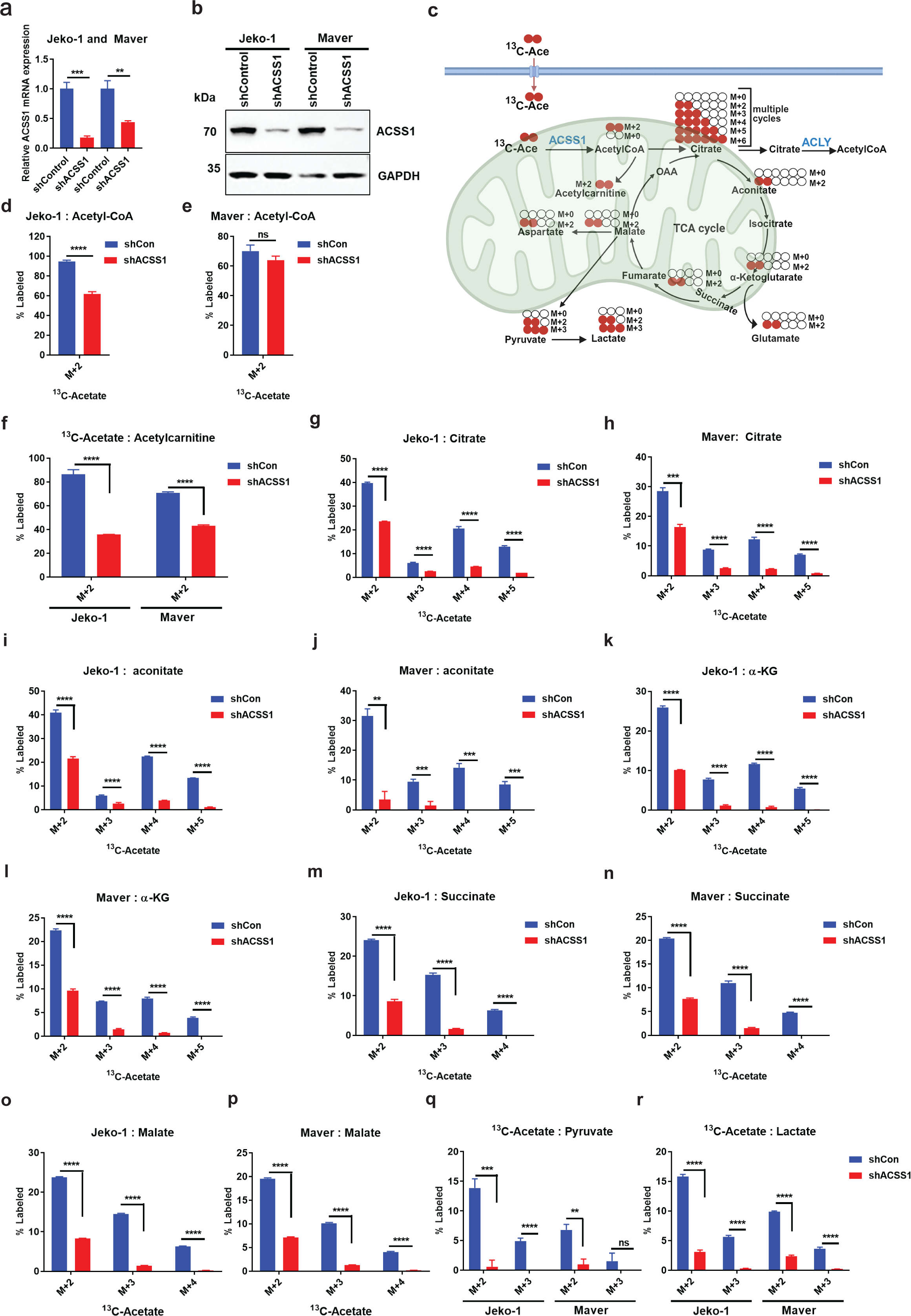
Mitochondrial ACSS1 knockdown impaired ^13^C-acetate mass isotopologues distributions (MIDs) in mantle cell lymphoma cell lines. **a,** q-PCR analysis of ACSS1 expression in scramble control (shCon) and shACSS1 knockdown (ACSS1 KD) in Jeko-1 and Maver cell lines. **b**, Immunoblotting analysis to confirm ACSS1 protein expression in Jeko-1 and Maver cell lines. **c,** schematic of labeled carbon molecules from ^13^C-acetate in multiple rounds of the TCA cycle is shown in red. **d-e,** Acetyl-CoA enrichment in shCon and shACSS1 Jeko-1 and Maver cells. **f,** Acetylcarnitine enrichment in shCon and shACSS1 cells. **g-h,** citrate enrichment in shCon and shACSS1 Jeko-1 and Maver cells. **i-j,** aconitate enrichment in shCon and shACSS1 Jeko-1 and Maver cells. **k-l**, α-KG enrichment in shCon and shACSS1 Jeko-1 and Maver cells. **m-n**, succinate enrichment in shCon and shACSS1 cells. **o-p,** malate enrichment in shCon and shACSS1 cells. **q-r,** pyruvate, and lactate enrichment in shCon and shACSS1 cells. All graphs show mean ± SD (n = 3 biological replicates), and all statistical analyses were conducted with unpaired t-tests: ^∗^p < 0.05, ^∗∗^p < 0.01, ^∗∗∗^p < 0.001, ^∗∗∗∗^p < 0.0001.

### 13C-acetate tracing analysis reveals Mitochondrial ACSS1 role in *de novo* pyrimidine metabolism

Based on the ^13^C-acetate tracing data, we found a noticeable difference in metabolizing the ^13^C-acetate in the TCA cycle intermediates between the ACSS1 KD and the control cells. To assess the broader impact on metabolic pathways, we examined the overall levels of steady-state metabolites in Jeko-1 to identify significant changes between ACSS1 KD and control cells. Out of the 148 detected metabolites, 48 showed a statistically significant fold change (>1.5 up or down, p<0.05). The findings revealed downregulation of *de novo* pyrimidine nucleotide synthesis pathway metabolites in ACSS1 KD cell lines (Extended Data Fig. 4a). Similar effects on metabolite abundance were observed in Maver cell lines. However, the effects were less pronounced than in Jeko-1 (Extended Data Fig. 4b). Our metabolomics analysis showed that ACSS1 KD was associated with reduced steady-state levels of metabolites in the pyrimidine metabolism pathway. To determine the contribution of acetate to this pathway, we examined the precursors for pyrimidine synthesis pathways in the ^13^C-acetate isotope tracing data as shown in the schematics (Fig. 4a). The results showed a highly significant reduction in ^13^C-acetate derived M+2-5 carbon labeling pattern into Glutamate (M+2-5; P<0.0001) in ACSS1 KD Jeko-1 and Maver cell lines compared to control (Fig. 4b,c). Next, we assessed the labeling of aspartate, and the results revealed a significant reduction in ^13^C-acetate derived aspartate (M+2-5; P<0.0005 and M+2-5; P<0.001) MIDs in Jeko-1 and Maver ACSS1 KD compared to control, respectively (Fig. 4d,e). Our findings emphasize the importance of ACSS1 in providing two essential precursors, glutamate and aspartate, for de novo pyrimidine synthesis and in promoting cell proliferation through a series of enzymes, including multienzyme complex CAD (carbamoyl phosphate synthetase), aspartate carbamoyltransferase (ATC) and dihydroorotate dehydrogenase (DHODH). The results showed a significant decrease in the labeling of DHO M+2 isotopologue in ACSS1 KD Jeko-1 and Maver cells compared to control cells (Jeko-1, p < 0.0001, Maver, p < 0.00001) (Fig. 4f,g). Additionally, we investigated the conversion of DHO substrate to orotate (OA) and the carbon labeling pattern from the ^13^C-acetate carbon source. The results showed that reduced DHO to orotate conversion in ACSS1 KD cells compared to controls, with a drastic reduction in % labeling of M+2 carbons (Jeko-1, p < 0.001, Maver, p < 0.0001) (Fig. 4hi). The significantly changed pyrimidine pathway metabolites were shown in the heatmap (Fig. 4j). To investigate the impact on metabolic pathways further, we used the web-based software MetaboAnalyst 5.0 for the enrichment and pathway analysis on ACSS1 KD and controlled MCL cell lines^34,43^. The pathway impact and enrichment analysis revealed that the pathways of pyrimidine metabolism as the top pathway, along with other pathways such as the citrate cycle (TCA cycle) and Alanine, Aspartate, and Glutamine amino acids were affected in ACSS1 KD (Fig. 4k, l). We hypothesized that overexpressing ACSS2 could compensate for the loss of ACSS1. To further investigate, we cultured Maver control and ACSS1 KD cell lines under nutrient-deprived conditions and performed qPCR analysis. The data revealed a more than 5-fold increase (P < 0.006) and an 8.0-fold increase (P < 0.01) in ACSS1 expression in control cells cultured in glucose-free (Glc) and 2mM acetate media, respectively, compared to cells cultured in complete RPMI with 25 mM glucose, 10% FBS, and 2 mM glutamine media (Fig. 4m). We also observed a more than 3-fold increase (P < 0.001) in ACSS1 expression in ACSS1 KD cells cultured in similar conditions. Further, the results revealed an increase of more than 2-fold, 3.0-fold and 5-fold ACSS2 expression in ACSS1 KD cells as a compensatory mechanism when cultured in nutrient-deprived conditions (Fig.4n).

**Fig. 4.**
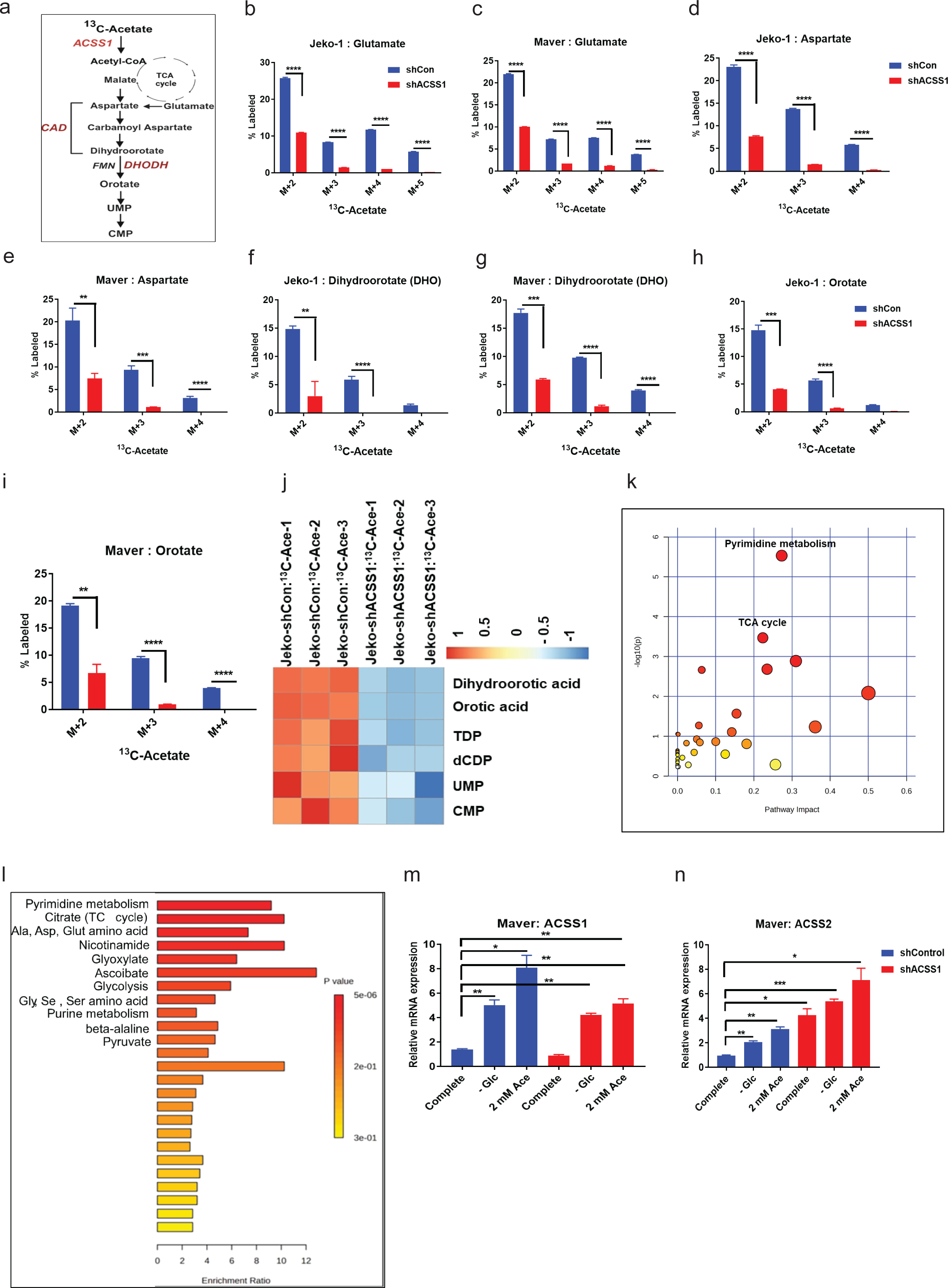
Mitochondrial ACSS1 regulates De Novo Pyrimidine biosynthesis in mantle cell lymphoma cell lines. **a,** A schematic of ^13^C-acetate tracing and mass isotopologues distributions in pyrimidine pathway metabolites. **b-c,** ^13^C-acetate derived M+2-5 labeled glutamate in Jeko-1 and Maver shCon and shACSS1 cells. **d-e,** ^13^C-acetate derived M+2-4 labeled aspartate in Jeko-1 and Maver shCon and shACSS1 cells. **f-g,** ^13^C-acetate derived M+2-4 labeled Dihydroorotate (DHO) Jeko-1 and Maver shCon and shACSS1 cells. **h-i,** ^13^C-acetate derived M+2-4 labeled orotate Jeko-1 and Maver shCon and shACSS1 cells. **j,** The heatmap shows a significantly changed metabolite in shCon vs. shACSS1 in Jeko-1 cell lines linked to *de novo* pyrimidine synthesis. **k-l**, The pathway impact and enrichment analysis. **m-n**, The q-PCR analysis of ACSS1 and ACSS2 expression in Maver, shCon, and shACSS1 cells cultured in various nutrient-deprived conditions. All graphs show mean ± SD (n = 3 biological replicates), and all statistical analyses were conducted with unpaired t-tests: ^∗^p < 0.05, ^∗∗^p < 0.01, ^∗∗∗^p < 0.001, ^∗∗∗∗^p < 0.0001.

### ACSS1 regulates oncometabolite 2-hydroxyglutarate (2HG) synthesis

^13^C-isotope tracing analysis using glucose and glutamine on RL, Jeko-1, and Maver cell lines revealed no significant differences in the fraction of M+2 labeled 2HG in RL compared to Jeko-1 and Maver cell lines (Fig.5 a, b). Next, in ^13^C-isotope tracing analysis on Jeko-1, Maver, and RL cell lines revealed significant differences in the fraction of M+2 labeled 2HG derived from ^13^C-acetate. (RL vs. Jeko-1, p<0.0001; RL vs. Maver, p<0.0001) compared to ^13^C-glucose and ^13^C-glutamine (Fig.5 c). We focused on the contribution of acetate to 2HG synthesis by performing ^13^C-acetate tracing on ACSS1 KD and control Jeko-1 and Maver cells as depicted in the schematic (Fig.5 d). To investigate the role of ACSS1 in acetate-derived 2HG isotopologues labeling, we used control and ACSS1 KD Jeko1-1 and Maver cell lines. The data revealed that a significant decrease in ^13^C-acetate derived 2HG in Jeko-1 (con, >26%; shACSS1 > 8%; p<0.0001) and Maver (% con vs shACSS1 M+2, -68%; M+3 -90% ; p<0.0001) cells with ACSS1 KD compared to control cells (Fig. 5e, f). It’s been previously reported that an increased succinate/αKG ratio in cells drives histone demethylase activity and regulates histone methylation epigenetically^44^. We assessed the succinate/αKG ratio based on the peak abundance. The data revealed an elevated succinate/αKG ratio in Jeko-1 and Maver cell lines (Fig. 5g). Based on our reasoning, we hypothesized that overactive IDH1/2 enzyme might directly convert α-KG to 2HG. We examined the expressions of IDH1 and IDH2. We found that cytosolic IDH1 is differentially expressed in MCL cell lines, especially overexpressed in the IBR-resistant Maver cell line (Fig. 5h). Mitochondrial IDH2 is highly expressed in all three cell lines (Fig. 5i). Additionally, we examined the expression of Nuclear Receptor Binding SET Domain Protein 2 (NSD2), also known as WHSC1, due to its role in histone methylation in the context of increased levels of 2HG^45^. It is also one of the common baseline mutated genes in MCL patients^46^. Next, we assessed the expression of WHSC1 in control and ACSS1 KD Jeko-1 cell lines by Q-PCR analysis. The results revealed decreased WHSC1 expression in the ACSS1 KD cell line compared to the control (Fig. 5j). The schematic diagram demonstrates how mitochondrial ACSS1 is involved in histone methylation (Fig. 5k). Based on increased succinate/α-KG and 2HG/α-KG ratios we developed a hypothesis regarding a potential new role for acetate metabolism in epigenetic regulation, specifically in histone methylation, in addition to its known involvement in histone acetylation. To investigate if histone methylation levels changed after ACSS1 KD, we cultured control and ACSS1 KD Jeko-1 and Maver cell lines in nutrient-deprived conditions. The results revealed that reducing ACSS1 led to significantly higher levels of the methylated H3K36me2 and H3K36me3 in cells grown in 2 mM acetate for 48 hours (Fig. 5l). Our research suggests that the increased succinate/α-KG ratios were associated with elevated histone methylation. In a previous study, the role of acetate-derived acetyl-CoA through ACSS2 on histone acetylation was investigated^17,47–49^. The role of ACSS1 in histone acetylation and epigenetic regulation has not been previously demonstrated. Our results showed that in nutrient-deprived conditions, H3K9 acetylation was decreased in ACSS1 KD cell lines cultured in 5 mM Glucose compared to shControl (Fig.5l). We investigated the impact of de novo pyrimidine synthesis and histone methylation by treating control and ACSS1 KD cell lines with the small molecule inhibitor of DHODH, brequinar and ACSS2 inhibitor, for 72 hours. After the treatment, we conducted a western blot analysis. The results indicated that brequinar enhanced the methylation of histone H3K27me3, H3K36me2, and H3K36me3 while decreasing histone acetylation in ACSS1 KD cells (Fig. 5m). However, the combination of ACSS2i+ brequinar treatment decreased the methylation of histone H3K27me3, H3K36me2, and H3K36me3 and H3K9 acetylation in ACSS1 KD cells (Fig. 5m). In contrast, under conditions where ACSS2 was inhibited, we observed decreases in both histone methylation and acetylation (Fig. 5m).

**Fig. 5.**
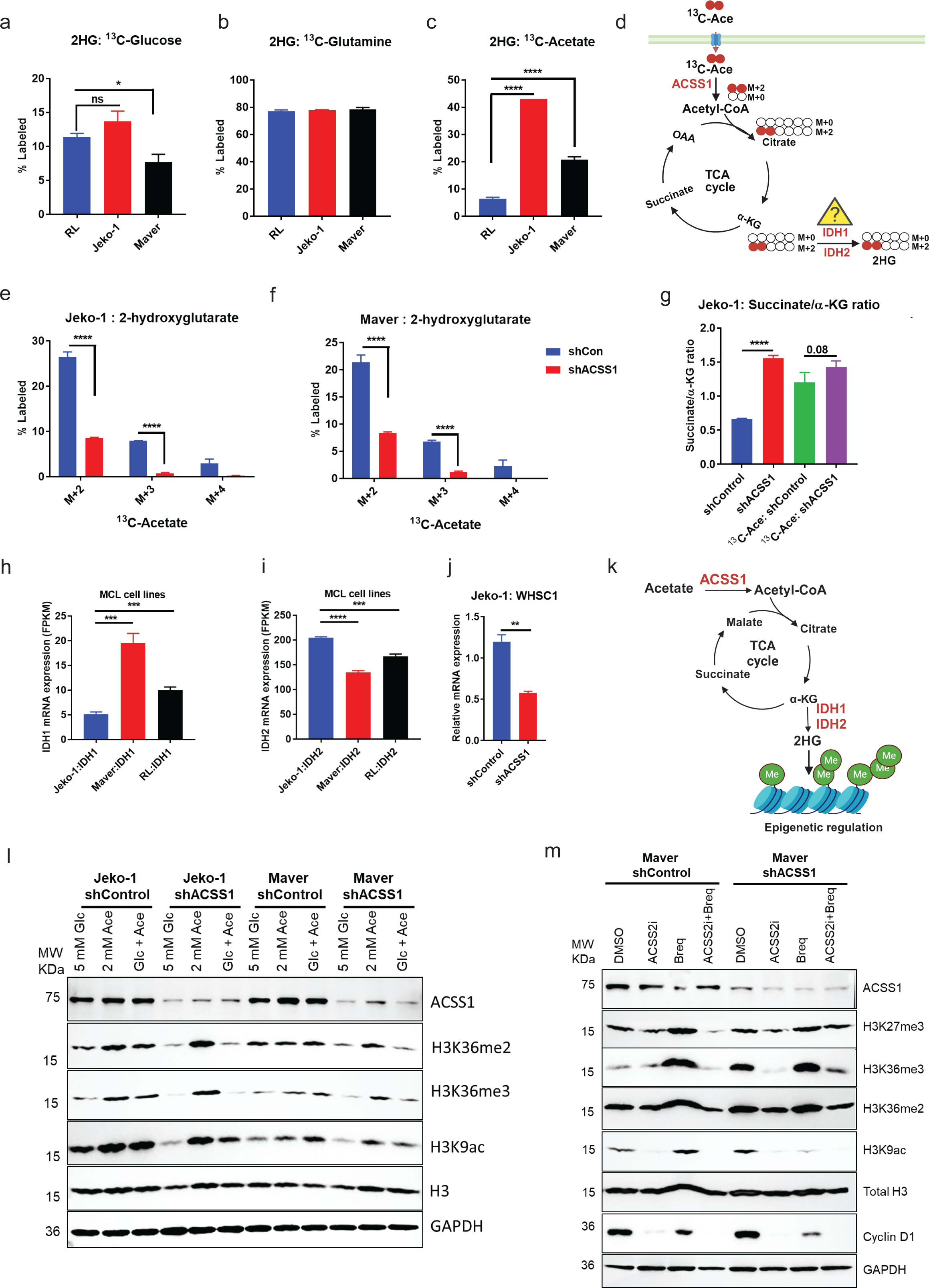
Mitochondrial ACSS1 converts acetate to oncometabolite 2-hydoxyglutarate (2HG) biosynthesis and regulates Histone methylation in mantle cell lymphoma cell lines. **a-c,** Differential % labeling pattern in 2HG from ^13^C-acetate, ^13^C-glucose, and ^13^C-glutamine-derived carbon sources is observed. **d,** A schematic of ^13^C-acetate derived M+2 labeled 2HG and glutamate carbon molecules is shown in red. **e-f,** Fractional enrichment of M+2-5 labeled 2HG derived from ^13^C-acetate in shCon and shACSS1 Jeko-1 and Maver cell lines. **g**, Succinate/ α-KG ratio, in Jeko-1 shCon and shACSS1 cells. **h-i,** RNA-seq analysis on MCL cell line expressing IDH1 and IDH2. **j**, q-PCR analysis on WHSC1 (NSD2) in shCon and shACSS1 Jeko-1 cell line. **k**, A schematic of ^13^C-acetate derived metabolites and enzymes linked Histone methylation. The genes confirmed in RNA-seq data is shown in red. **l**, Jeko-1 shCon and shACSS1 cells cultured in nutrient-deprived conditions. **m**, Jeko-1 parental cells were cultured in complete media in DMSO control or the presence of DHODH inhibitor, brequiner (10 uM) alone or ACSS2i (10 uM) in combination with brequiner for 48 hours, the lysates were probed for the indicated antibodies in immunoblotting. All graphs show mean ± SD (n = 3 biological replicates), and all statistical analyses were conducted with unpaired t-tests: ^∗^p < 0.05, ^∗∗^p < 0.01, ^∗∗∗^p < 0.001, ^∗∗∗∗^p < 0.0001.

### Mitochondrial ACSS1 regulates acetate metabolism and oxygen consumption (OCR) in nutrient-deprived conditions

To investigate the potential impact of ACSS1 on acetate-mediated OCR, we measured the basal OCR level in RL (low ACSS1 expressing), Jeko-1, and Maver (high ACSS1 expressing) cell lines by exposing glucose-starved cells to 2 mM acetate. The results showed that acetate increased OCR by 24% (p < 0.002) and 30% (p < 0.0001) in Jeko-1 and Maver cells as compared to RL cells, respectively. However, we did not observe a significant change in RL cells (Fig. 6a). These results suggest that ACSS1 is responsible for the increased acetate-fueled mitochondrial respiration. We then compared the OCR in control and ACSS1 KD Jeko-1 and Maver cell lines. The results showed a decrease in the basal and under-stressed condition OCR in the ACSS1 KD cells compared to the control Jeko-1 (Fig. 6b) and Maver cells (Fig. 6c). Next, we subjected the Jeko-1 and Maver parental cell lines to total, cytosol, and mitochondrial fractionations for immunoblotting and the results revealed that ACSS1 and DHODH were predominantly located in the mitochondria (Fig. 6d). In contrast, CAD was in the cytosol. Next, we examined cyclin D1, a significant oncogene in MCL pathogenesis. Our results showed that cyclin D1 was predominantly present in the cytosol and only in small quantities in the mitochondrial fraction. Based on the literature, DHODH is the only enzyme involved in *de novo* pyrimidine synthesis that is predominantly located in the mitochondria. However, a recent study has shown that DHODH is localized in mitochondria and the nucleus, implying its unknown functional role in addition to catalyzing dihydroorotate^50^. To investigate whether DHODH, which is involved in pyrimidine synthesis, can translocate to the nucleus, we performed cytosol/mitochondrial and nuclear fractionation on Jeko-1 and Maver cell lines treated with DMSO control and the DHODH inhibitor brequimer, followed by western blotting. The results indicated that DHODH was found in the nuclear fractionations (Fig 6e).

**Fig. 6.**
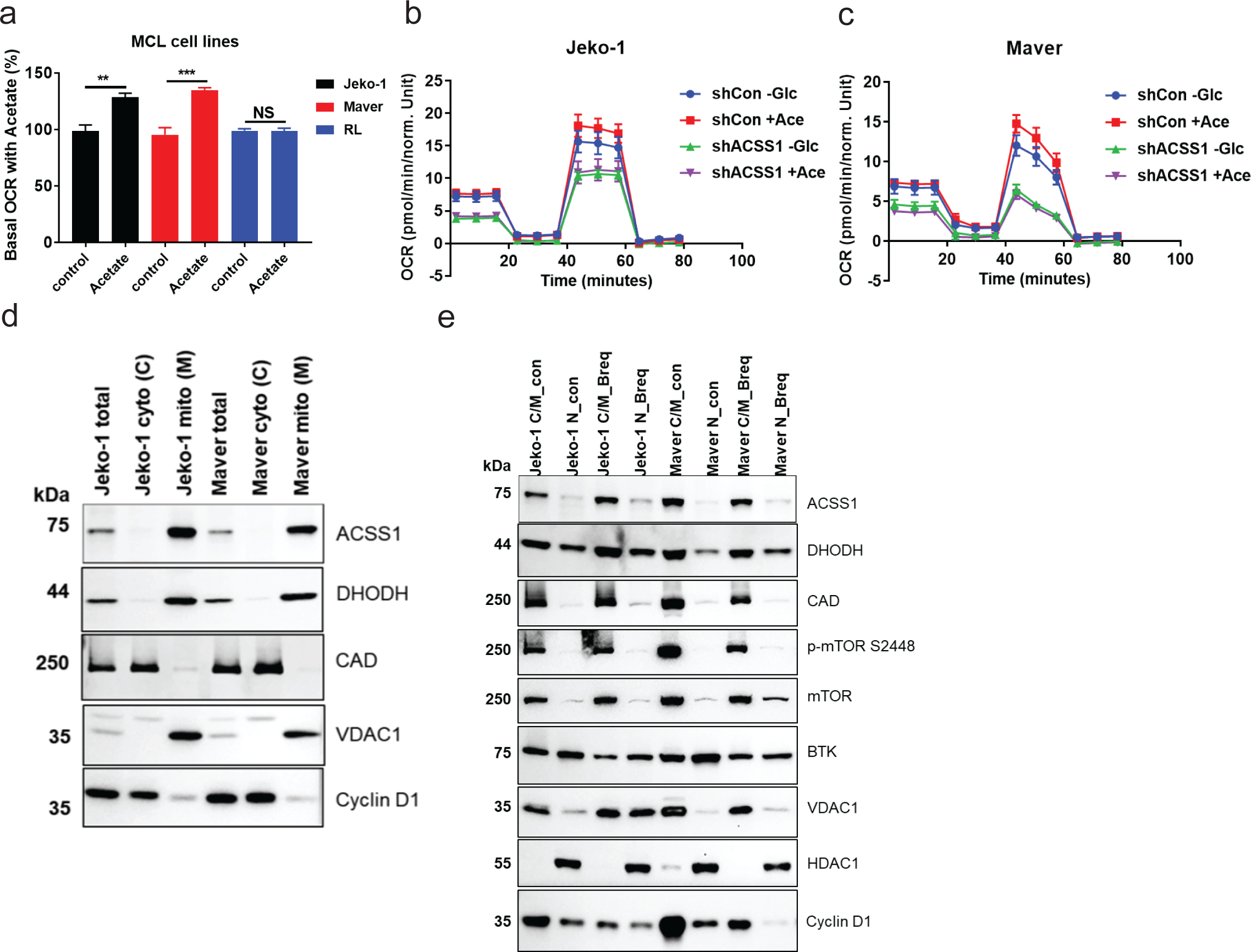
Mitochondrial ACSS1 regulates oxygen consumption in mantle cell lymphoma cell lines. **a,** Histogram of percent basal Oxygen consumption rate (OCR) in three MCL cell lines in ACSS1 high (Jeko-1 and Maver) and low (RL) expressing cell lines (N=5). **b-c,** The OCR measurement in shCon and shACSS1 Jeko-1 and Maver cell lines (N=7). **d,** Total, cytosol, and mitochondrial fractionation of Jeko-1 and Maver parental cell lines and probed with indicated antibodies. **e,** Cytosol/mitochondrial and nuclear fractionation of Jeko-1 and Maver parental cell lines treated with DMSO or DHODH inhibitor, brequiner (10 uM) for 24 hours and probed with indicated antibodies. All graphs show mean ± SD (> n = 3 biological replicates), and all statistical analyses were conducted with unpaired t-tests: ^∗^p < 0.05, ^∗∗^p < 0.01, ^∗∗∗^p < 0.001, ^∗∗∗∗^p < 0.0001.

### Mitochondrial ACSS1 is critical for MCL cell growth and viability in nutrient-deprived conditions

Earlier reports showed that glucose deprivation increased ACSS1 and ACSS2 expression in MEL697 melanoma cells and that ACSS1 or ACSS2 depletion reduced tumor growth^51^. Having established the role of ACSS1 in *de novo* pyrimidine synthesis, we reasoned that it could also play an essential role in cell growth and survival under nutrient-deprived conditions. We used ACSS1 KD and control cell lines. The efficacy of ACSS1 KD was validated through western blotting (Fig. 7a). In the Maver and Jeko-1 cell lines, two of the four shRNA ACSS1 clones showed nearly 100% knockdown of ACSS1. We monitored cell growth and viability using an automated cell counter (Thermofisher) and trypan blue staining. The data showed a significant decrease in cell growth (>70%; p<0.0001) and viability (>80%; p<0.0003) in the shRNA targeting ACSS1 group compared to the control in the Maver and Jeko-1 cell lines (Fig. 7b, c). We also evaluated the role of ACSS1 in the Jeko-1 cell line by comparing ACSS1 KD to the control and confirmed similar results to those observed in the Maver cell lines (Fig. 7d, e). Next, we cultured control and ACSS1 KD cell lines in complete media (RPMI with 25 mM glucose, 2 mM glutamine, 10% FBS, and 1% penicillin/streptomycin antibiotics), conditioned media (-Glc, -Gln, and 2 mM Acetate in 1% dialyzed FBS) and -Glc and -Gln media +1% dialyzed FBS for 72 hours. The results indicate no significant difference between the shCon and shACSS1 cells cultured under complete RPMI media in both Jeko-1 and Maver cell lines. However, when cultured under nutrient-deprived conditions, the results showed a significant reduction in cell viability in the ACSS1 KD cell lines compared to control. The acetate supplement increased cell viability in the control cell lines but did not have the same effect in the ACSS1 KD cell lines. This indicates that ACSS1 is crucial for metabolizing acetate to support cell survival under nutrient-deprived conditions. Next, we focused our efforts on delineating the effect of the ACSS2 inhibitor on the Jeko-1 and Maver parental cell growth by treating them with a small molecule inhibitor of ACSS2^22^. The results showed decreased cell growth of Jeko-1 and Maver in ACSS2i-treated cells in a dose-dependent manner (Fig.7f, g). Next, we assessed the cell death by annexin V staining by flow cytometry on Jeko-1 school and shACSS1 cell lines cultured in nutrient-deprived conditions. Acetate supplement rescued the cell death and decreased % of annexin V positive cells under nutrient-deprived conditions in control compared to the ACSS1 KD cell line (Fig. 7h, i). In our schematic, we summarize our findings about the role of ACSS1 under nutrient-deprived conditions. ACSS1 regulates the function of the TCA cycle by supplying all the TCA cycle intermediates. It also provides glutamine and aspartate precursors for synthesizing de novo pyrimidines for cell proliferation and D-HG oncometabolite for regulating histone modifications (Fig. 7j).

**Fig. 7.**
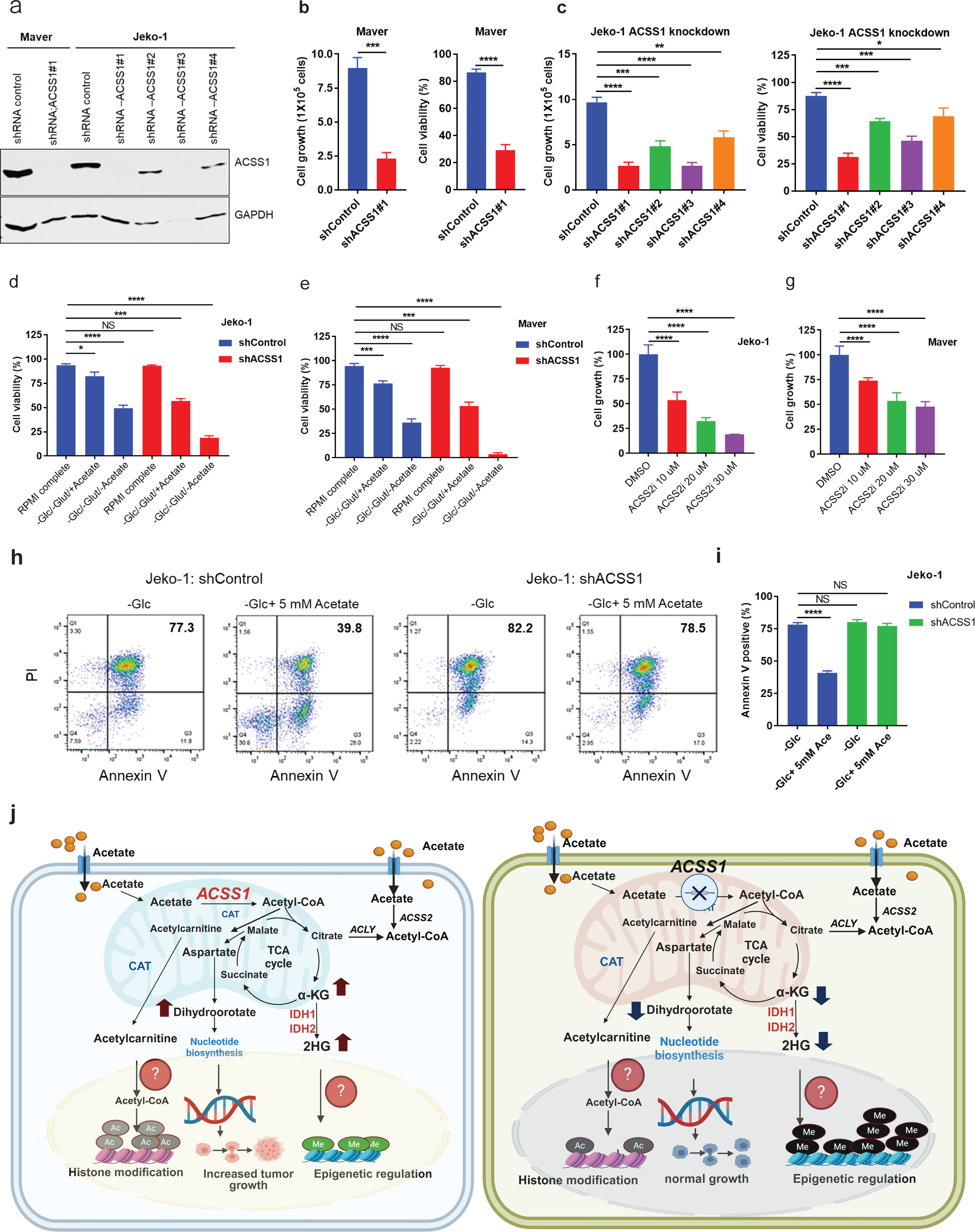
Mitochondrial ACSS1 regulates cell growth and viability under nutrient-deprived conditions in MCL. **a,** Immunoblotting confirms the shRNA-mediated knockdown of ACSS1 in MCL cell lines, probed with indicated antibody. **b-c**, ACSS1 knockdown impairs cell viability in Maver and Jeko-1 cell lines. **d-e,** ACSS1 knockdown cells cultured in nutrient-deprived conditions impair cell growth and viability in Maver and Jeko-1 cell lines. **f-g,** Jeko-1, and Maver parental cell lines treated with ACSS2i. **h,** Jeko-1 shCon and shACSS1 cultured in minus Glc and 5 mM acetate and PI and Annexin V staining for apoptosis assay. **i,** Histogram showing Annexin V positive cells (N=3). **j,** Graphical summary of the findings. ACSS1 converts acetate to acetyl-CoA and Acetylcarnitine for histone acetylation and utilizes glutamate, aspartate, and dihydroorotate for de novo pyrimidine synthesis. Additionally, ACSS1 converts acetate to the oncometabolite D-2HG and supports histone methylation for epigenetic regulation. All graphs show mean ± SD (n = 3 biological replicates), unless otherwise specified, and all statistical analyses were conducted with unpaired two-tailed t test: ^∗^p < 0.05, ^∗∗^p < 0.01, ^∗∗∗^p < 0.001, ^∗∗∗∗^p < 0.0001.

## Discussion

In our study, we discovered the crucial role of mitochondrial ACSS1 in acetate metabolism. We established its significant contribution to the conversion of the oncometabolite 2-hydroxyglutarate and demonstrated its essential involvement in *de novo* pyrimidine synthesis for the first time. These findings enhance our understanding of key metabolic pathways and their implications in cellular function. In mammals, acetyl-CoA is a central metabolite derived from glucose, glutamine, and fatty acids. Acetate can also be converted to acetyl-CoA by cytosolic ACSS2. Previous studies have reported an increase in acetate metabolism in multiple types of cancers, including prostate, liver, lung, brain, and breast cancer^23,52–55^. We discovered a previously unknown mechanism for the synthesis of the acetate-derived oncometabolite 2-hydroxyglutarate under nutrient-deprived conditions and its effect on histone methylation patterns through ACSS1.

We have demonstrated that ACSS1 is highly expressed in hematological and solid tumor cancer models by analyzing the publicly available TCGA database. Furthermore, we validated this finding by studying MCL, CLL, and DLBCL cases. Our data highlights the association between overexpression of ACSS1 and decreased sensitivity to ibrutinib, a small molecule inhibitor of Bruton’s tyrosine kinase (BTK). In our study, we observed that cell lines sensitive to ibrutinib (MCL-RL and OCI-LY1 in DLBCL) have lower ACSS1 expression compared to cell lines that are less sensitive or resistant to ibrutinib (Jeko-1, Maver, and Grant in MCL, and LY8 and Toledo in DLBCL). These findings indicate that ACSS1 plays a significant role in acetate-mediated OXPHOS and ibrutinib resistance in MCL and DLBCL cell lines. We utilized three ^13^C-labeled carbon sources: glucose, glutamine, and acetate, to determine the preferences for carbon sources in MCL cell lines expressing low and high levels of ACSS1. This method helped us to understand which energy sources MCL cell lines prefer for their metabolic processes in environments lacking nutrients. In normal or tumor conditions, cells predominantly use glucose and glutamine to create new biological molecules and produce ATP in the TCA cycle by generating TCA intermediates. We found that when we used ^13^C-labeled carbon sources to analyze the flux of carbon in MCL cells, we didn’t observe any significant difference in the use of glucose and glutamine between low and high ACSS1-expressing RL, Jeko-1, and Maver cell lines. However, when we looked at the flux of ^13^C-acetate, we did find a significant difference in the labeling pattern of TCA cycle intermediates in Jeko-1 and Maver cells with high ACSS1 expression (Fig. 2). This suggests that high ACSS1 expression allows MCL cell lines to survive under nutrient-deprived conditions by switching from glucose to acetate as an energy source. To further MCL cells with ACSS1 high expression, Jeko-1 and Maver cell lines and shRNA mediated knockdown ACSS1 study on ^13^C-acetate flux analysis demonstrated reduced ^13^C-acetate derived M+2-n carbon labeling pattern across TCA cycle intermediates, citrate, aconitate, αKG, succinate and malate (Fig.3). The metabolomics-based mass isotopologue distribution (MIDs) study showed that in ACSS1 overexpressing Jeko-1 and Maver MCL cell lines, acetate is used as a nutritional source in an ACSS1-dependent manner, more so than glucose or glutamine. This leads to acetate-mediated bioenergetics and cancer progression. Moreover, when glucose is deprived, ACSS1 overexpressing cell lines use acetate to produce TCA intermediates for OXPHOS and convert pyruvate and lactate for biomass growth through gluconeogenesis pathways (Fig. 3). In normal physiology, glucose is converted into pyruvate. The pyruvate then enters the TCA cycle to provide an efficient energy source. In mantle cell lymphoma cell lines experiencing nutrient deprivation, mitochondrial acetate is crucial in supplying acetyl-CoA through ACSS1. This promotes the efficient operation of the TCA cycle by providing all the necessary intermediates for ATP generation, histone modification via acetyl-CoA/acetylcarnitine for acetylation, and glutamate and aspartate precursors for the synthesis of nucleotides. The most important aspect of our finding is the conversion of acetate to 2-hydroxyglutarate and its effects on histone methylation. There are two pathways involved in pyrimidine synthesis in mammals: 1) the salvage pathway, which is active in resting or fully differentiated cells, and 2) the *de novo* pathway, which is active in highly proliferating cells^56^. Almost 40 years ago, Aitken and Lipmann discovered that ^14^C-acetate was incorporated into pyrimidine and DNA in the MCF-7 human breast cancer cell line, regulated by estrogen hormone^31^. The report did not identify any specific metabolic enzymes or mechanisms. Therefore, we are the first to report that the crucial mitochondrial acetate metabolic enzyme ACSS1 produces the essential precursor for pyrimidine synthesis. We demonstrated the role of ACSS1 in Jeko-1 and Maver cell lines using the ACSS1 knockdown strategy to support the hypothesis that ^13^C-acetate incorporates into pyrimidine. Our ^13^C-acetate tracing data under ACSS1 shControl and shACSS1 KD Jeko-1 and Maver cells reveal decreased M+2-5 glutamate and M+2-4 aspartate labeling precursors for nucleotide synthesis. Thus, our study shows that new role of ACSS1 in mitochondrial acetate-derived glutamate and aspartate carbon in pyrimidine nucleotides and their derivatives (i.e., DHO, orotate). The expression of ACSS1 is notably high in various cancer types, yet its role in cancer progression remains underexplored. Our data indicates that ACSS1 is involved in acetate metabolism and exhibits differences across MCL cell lines. It demonstrates a preference for utilizing acetate over glucose or glutamine in both low- and high-ACSS1-expressing cell lines. Notably, when comparing carbon sources from ^13^C-labeled glucose, glutamine, and acetate in TCA cycle intermediates, acetate is clearly a prominent carbon source. These findings suggest that ACSS1 is overexpressed in MCL cell lines and exhibits high enzymatic activity. Additionally, an enzymatic study of ACSS1 revealed that the Km value for acetate as a substrate is approximately 10 mM, compared to 0.1 mM for the cytoplasmic enzyme ACSS2. This represents a nearly 100-fold difference in substrate specificity for acetate^57^. Based on an earlier report on an enzymological study *in vitro* enzyme activity in rat liver shows that the Km value of the ACSS1 for acetate is approximately 10 mM when compared with approximately 0.1 mM for the cytoplasmic enzyme ACSS2, which is almost 100 fold difference in substrate specificity for acetate^58,59^The metabolomics-based mass isotopologue distribution (MIDs) study showed that in ACSS1-overexpressing Jeko-1 and Maver MCL cell lines, acetate is used as a nutritional source in an ACSS1-dependent manner, more so than glucose or glutamine. This leads to acetate-mediated bioenergetics and cancer progression.

Our results demonstrate that Jeko-1 and Maver cells are viable in glucose-free media replaced with 1 to 2 mM sodium acetate. Cancer cells have upregulated glycolysis compared with normal cells. Recent studies have shown that OXPHOS can also be upregulated in certain cancers, including leukemia, lymphomas, pancreatic ductal adenocarcinoma, high OXPHOS subtype melanoma, and endometrial carcinoma^60^. However, in specific melanoma cells, glucose-independent acetate metabolism promotes tumor growth in mice^51^. Therefore, ^11^C-acetate positron emission tomography (PET) positivity has been used in the diagnosis of several cancer models, prostate^61^, gliomas^62^ and increased ^11^C-acetate accumulation in IDH-mutated human glioblastoma^63^ and it is linked to ACSS2-mediated acetate metabolism. Based on the previous study, ^11^C-acetate accumulation in primary prostate tumors was positive in all patients, and (11) C-acetate PET in a patient with lymph node metastasis showed high intrapelvic accumulation corresponding to metastatic sites, suggesting (11) C-acetate PET can be used for the diagnosis of MCL and DLBCL patients.

Our findings suggest that metabolites derived from acetate in the tricarboxylic acid (TCA) cycle can enhance oxidative phosphorylation (OXPHOS) and promote the production of the oncometabolite 2-hydroxyglutarate. This mechanism may significantly influence epigenetic regulation through histone methylation in cancer models, particularly in mantle cell lymphoma and diffuse large B-cell lymphoma (DLBCL), where levels of ACSS1 are notably high under conditions of limited nutrient availability. Therefore, targeting mitochondrial ACSS1 with small molecule inhibitors could present a promising and innovative therapeutic strategy worth further exploration.

## Acknowledgments

We are grateful to Dr. Erica Golemis, Senior Associate Dean of Research at Lewis Katz School of Medicine, Temple University Health System, and Professor and Chair, Cancer and Cellular Biology, Fox Chase Cancer Center, Philadelphia, for critically reading the manuscript and the feedback to improve the paper. The Wistar Institute Proteomics and Metabolomics Shared Resource is supported by NIH Cancer Center Support Grant CA010815. The metabolomics analysis was performed on a Thermo Q-Exactive HF-X mass spectrometer purchased with NIH grant S10 OD023586. The Fox Chase Cancer Center startup funding for MW. We created the figures in this manuscript using the Biorender program.

## Author Contributions

J.B. conceived the project, designed and performed the research, analyzed the data, supervised the project, and wrote the manuscript; A.R.G., metabolomics analysis and manuscript review; C.L., heat map analysis, and review; S.W., D.R., O.M., N.V.S., V.S.M., P.J., H.Y., P.L., performed experiments; M.E., K.Q.C., R.N., performed IHC; H.B., resources; K.E.W., resources and manuscript review; M.A.W., resources, manuscript reviewing and acquiring the funding for the project. All authors read and approved the manuscript.

## Figure legends

**Extended Data Fig. 1.**
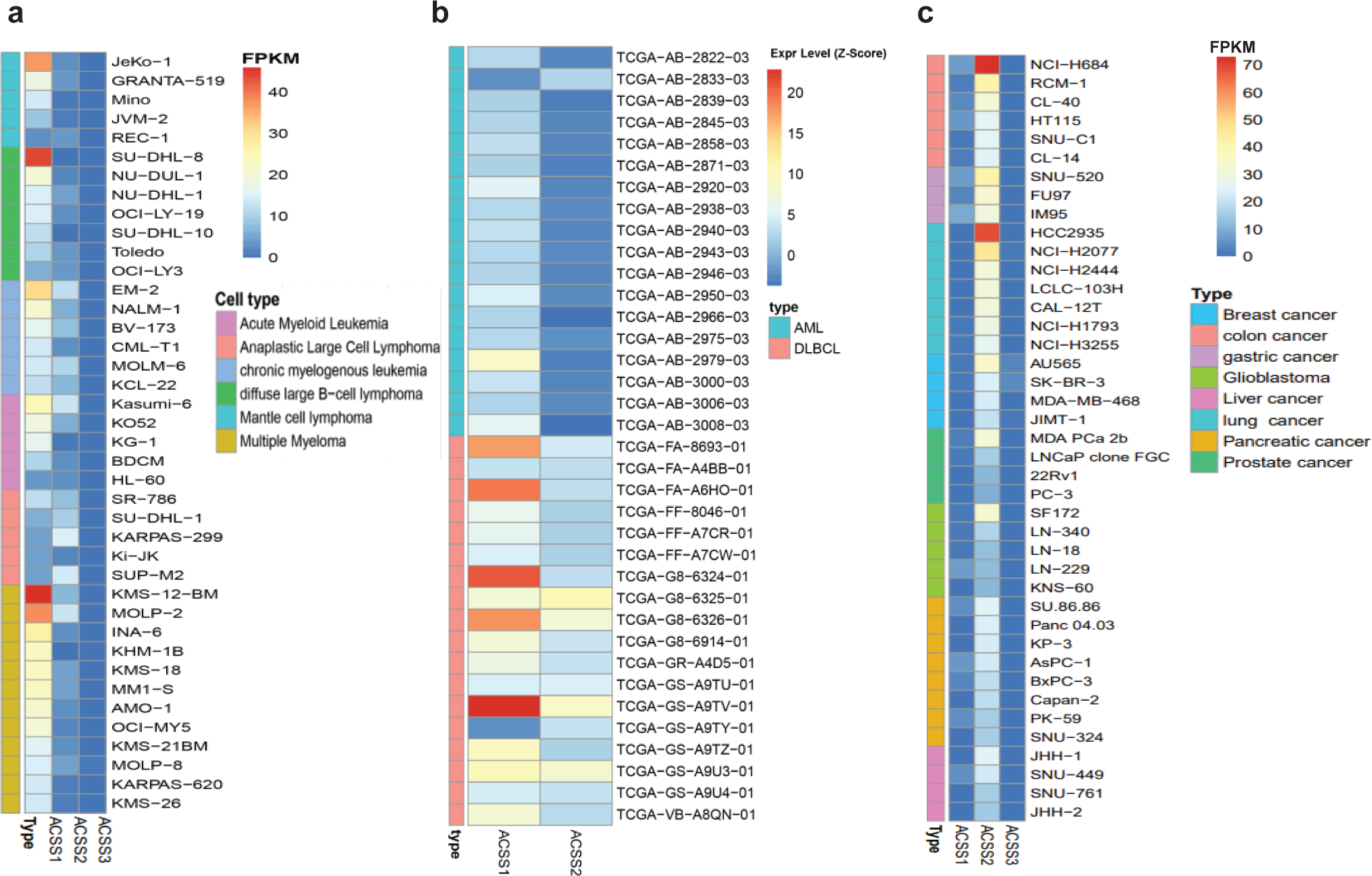
Mitochondrial acetate metabolic enzyme ACSS1 is overexpressed in hematopoietic malignancy cell lines and patient-derived samples. **a,** Bioinformatics analysis of selected gene expression in the cell line atlas database (https://www.ebi.ac.uk/gxa/home) revealed mitochondrial ACSS1 overexpression in hematopoietic relevant cell lines (heat map); **b**, ACSS2 is overexpressed in solid tumors (heat map) and **c**, The Cancer Genome Atlas (TCGA) database revealed ACSS1 overexpression in hematopoietic malignancy of AML and DLBCL patient samples.

**Extended Data Fig. 2.**
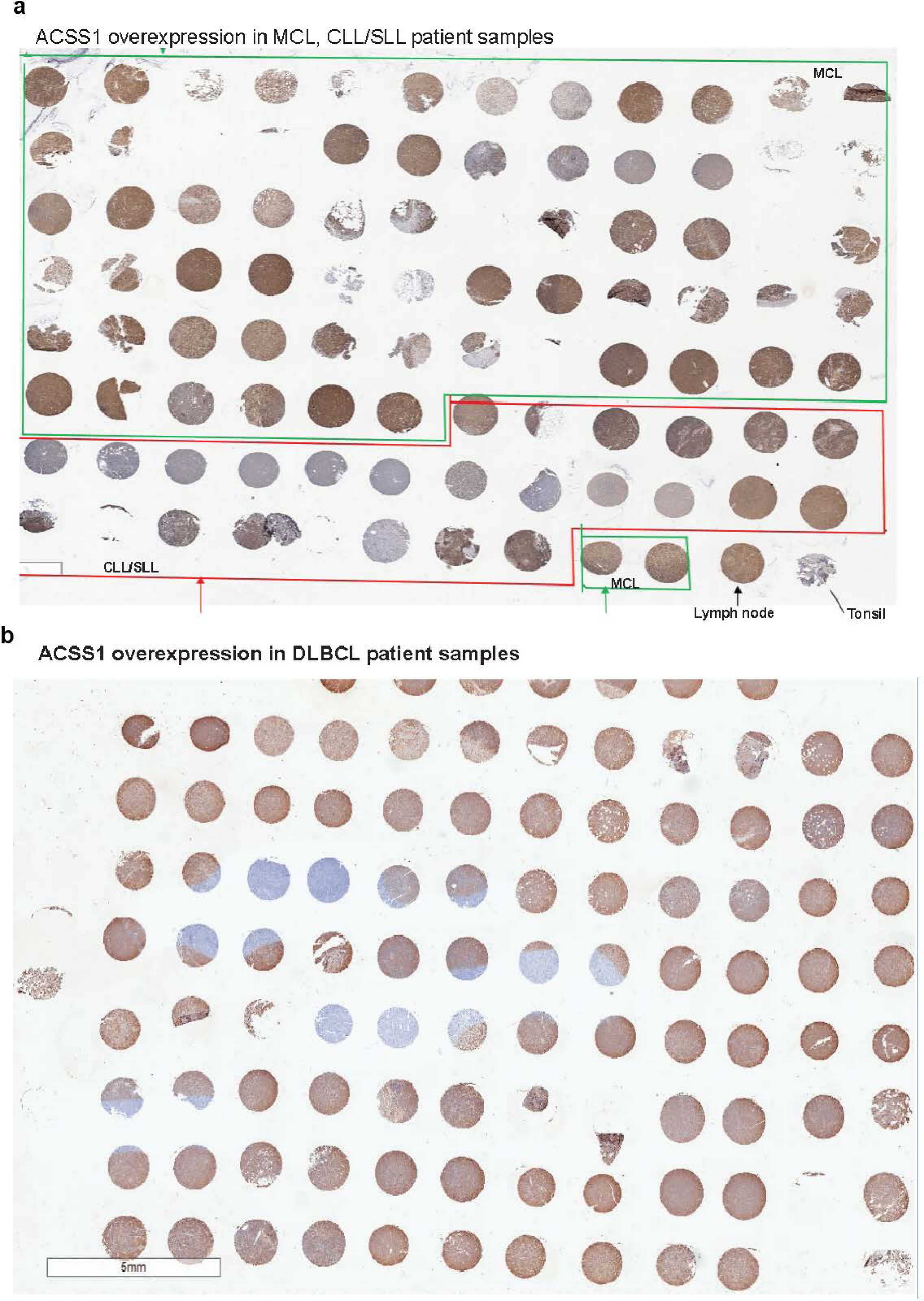
Immunohistochemistry (IHC) staining of ACSS1 in MCL, DLBCL, and ALCL patient-derived TMA blocks. **a,** ACSS1 overexpression in MCL, CLL/SLL patient samples; **b**, ACSS1 overexpression in DLBCL patient samples.

**Extended Data Fig. 3.**
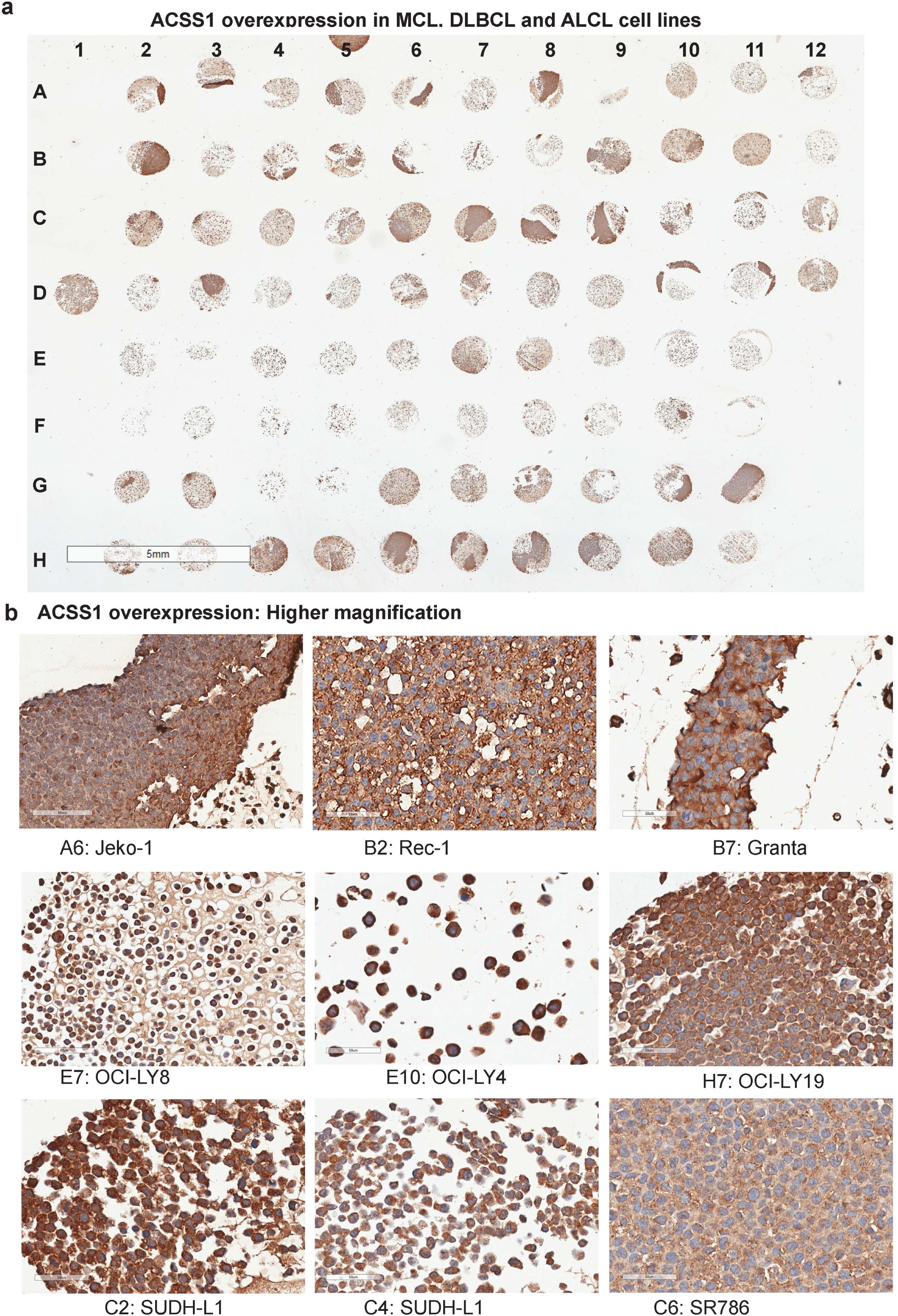
Immunohistochemistry (IHC) staining of ACSS1 in MCL, DLBCL, and ALCL cell line blocks. **a** ACSS1 overexpression in MCL, DLBCL and ALCL cell line derived blocks; b, ACSS1 overexpression in images in higher magnification from the cell line blocks, **top**; MCL cell lines, Jeko-1, Rec-1, and Granta; **middle**, DLBCL cell lines, OCI-LY8, OCI-LY4 and OCI-Ly19 and **bottom**, ALCL cell lines, SUDHL-1, SUPM2, and SR786.

**Extended Data Fig. 4.**
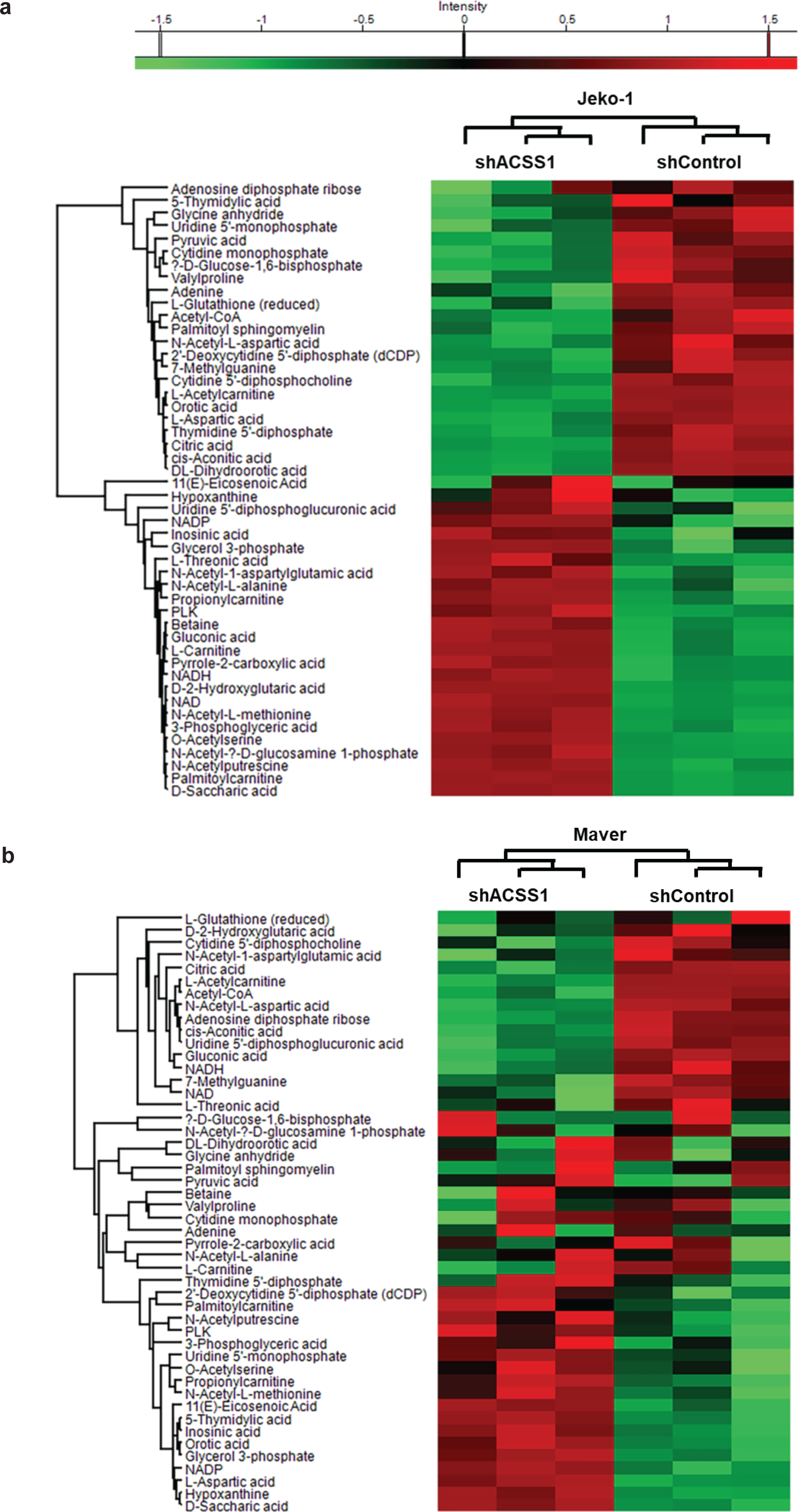
Mitochondrial ACSS1 and acetate metabolism revealed enrichment of de novo pyrimidine pathway metabolites. **a-b,** The Heatmap shows a significantly changed metabolite in shCon vs. shACSS1 in Jeko-1 and Maver cell lines. c-d, Pathway impact and enrichment analysis. e, The Heatmap shows a significantly changed metabolite-linked pyrimidine pathway in shCon vs. shACSS1 in the Jeko-1 cell line.

